# Towards a greener AlphaFold2 protocol for antibody-antigen modeling: Insights from CAPRI Round 55

**DOI:** 10.1101/2024.10.07.616947

**Authors:** Büşra Savaş, İrem Yılmazbilek, Atakan Özsan, Ezgi Karaca

## Abstract

In the 55^th^ round of CAPRI, we used enhanced AlphaFold2 (AF2) sampling and data-driven docking. Our AF2 protocol relies on Wallner’s massive sampling approach, which combines different AF2 versions and sampling parameters to produce thousands of models per target. For T231 (an antibody peptide complex) and T232 (PP2A:TIPRL complex), we employed a 50-fold reduced MinnieFold sampling and a custom ranking approach, leading to a top-ranking medium prediction in both cases. For T233 and T234 (two antibody bound MHC I complexes), we followed data-driven docking, which did not lead to an acceptable model. Our post-CAPRI55 analysis showed that if we would have used our MinnieFold approach on T233 and T234, we could have submitted a medium-quality model for T233 as well. In the scoring challenge, we utilized the scoring function of FoldX, which was effective in selecting acceptable models for T231 and medium quality models for T232. Our success, especially in predicting and ranking a medium quality model for T231 and potentially for T233 underscores the feasibility of green and accurate enhanced AF2 sampling in antibody complex prediction.

## INTRODUCTION

In the CASP14 competition, AlphaFold2 (AF2) achieved an unprecedented success rate in producing highly accurate protein structural models [1]. The release of AF2’s source code motivated the community to adapt AF2’s use in multimer modeling. In response to this interest, DeepMind extended the training regime of AF2 to support protein multimer modeling (leading to AF v2.1) [2]. The interfacial clash issues produced by AF v2.1 was resolved in the next 2.2 version. By employing these versions, the community achieved an extraordinary 90% success rate in multimer modeling in CASP15-CAPRI [3]. In this round, two teams achieved top-ranking predictions by using both AF v2.1 and v2.2 model weights. Among these two, the Venclovas team, played with input multiple sequence alignments (MSAs) and increased the number of recycles during AF2 sampling [4]. Wallner Team, on the other hand, kept the standard MSA generation protocol of AF2, but they played with several AF2 parameters to generate 6,000 models per target. For this, they modified AF2’s dropout, template use, number of recycles and number of seeds [5, 6]. This massive sampling strategy, as they called, AFSample, led to the submission of the highest number of high-quality model submissions [7].

Inspired by the success of AFSample, DeepMind released the third version of AFv2, which is retrained on a larger set of complexes, with increased number of recycles and seeds [8]. Expanding on these developments, in CAPRI55, we wanted to explore whether a reduced version of Wallner’s massive sampling strategy could produce meaningful results. Accordingly, we devised a 50-fold reduced version of this idea and called our approach as MinnieFold, since it is using 95% reduced GPU hours compared to a full massive sampling run with Wallner’s setup in CASP15-CAPRI [5]. MinnieFold was used to predict the binding of two targets (T231-2) during the competition and the other two (T233-4) after the competition. As an outcome, MinnieFold led to the generation of medium quality models for T231, T232, and T233. During the competition, we used data-driven docking for T233 and T234, which unfortunately did not result in an acceptable model. In the scoring round, we ranked the model pool with the interface scoring of FoldX, which selected acceptable and medium quality models for T231 and T232 targets, respectively [9].

In the following sections, we provide a detailed analysis of our performance in CAPRI55, highlighting our strategies, specifically in modeling antibody-antigen targets with MinnieFold (T231, T233-4), and their outcomes.

## METHODS

### Prediction Category

The summary of our modeling strategies is illustrated in Figure S1.

- Throughout the 55^th^ round, the ColabFold web resource was accessed through Colab Notebook version 1.5.3, post-CAPRI models were obtained with version 1.5.5 [10]: https://colab.research.google.com/github/sokrypton/ColabFold/blob/main/AlphaFold2.ipynb In the case of T232, we followed AFsample pipeline for scenarios involving v2.1 and v2.2 on a lab-scale workstation equipped with an A4000 GPU (https://github.com/bjornwallner/alphafoldv2.2.0).
- AF2’s refinement protocol: Amber relaxation was applied with max_iterations 2000, tolerance 2.39, stiffness 10.0. (https://colab.research.google.com/github/sokrypton/ColabFold/blob/main/beta/relax_amber.ipynb)
- AF3 was run on https://alphafoldserver.com/ [11]
- HADDOCK was accessed via https://rascar.science.uu.nl/haddock2.4/submit/1 [12]. In T232 case, the catalytic subunit of PP2A and TIPRL were merged into a single unit to follow a two-body docking with the following parameters: center of mass restraints=on, it0 = 5000, it1 = 400, and water = 400. For T233 and T234, each molecule was treated as a single unit, following four-body docking using the same sampling parameters.
- HADDOCK refinement protocol was accessed via https://rascar.science.uu.nl/haddock2.4/refinement/1 [12]. This protocol allows the refinement of the interface with position restraints applied to backbone atoms of the protein complex, excluding the interface atoms.
- AF2Rank is accessed via a Colab Notebook, introduced in https://github.com/jproney/AF2Rank [13]. While running AF2rank, we enabled amino acid and sidechain unmasking and used four recycles to ensure accurate reproduction of the input.
- The molecular dynamics (MD) simulations were run using GROMACS (2020.4) with CHARMM forcefield [14, 15].
- The contacts and interface energies are calculated with our in-house interface analysis tool, DynaBench [16].
- The Cα root mean square deviations (RMSD) are calculated with the `MULTI` command of ProFitV3.1, which is an implementation of the McLachlan algorithm [17].

We provide our MinnieFold pipeline through a GitHub repository containing all our CAPRI and post-CAPRI models, along with their analysis files (https://github.com/CSB-KaracaLab/MinnieFold).

### Post-CAPRI analysis

All structural illustrations on this paper were created PyMOL 2.5.2 [21]. For further evaluation of our models, we used the DockQ metric (https://github.com/bjornwallner/DockQ) [20]. We followed the standard CAPRI protocol for DockQ calculation.

## RESULTS

### T231: Our MinnieFold sampling combined with peptide pLDDT ranking was our key highlight in CAPRI55

Target T231 was an antibody bound to an eight amino acids long peptide (PDB ID: 8RMO [21]). Since this target lacked the typical co-evolution signal, we decided to use an enhanced AF2 sampling. For this, we constructed a mini version of Wallner’s AFsample protocol. In the regular AFsample, Wallner generates 1000 models per sampling scenario [6]. In our mini version, MinnieFold, we reduced this number by 50-fold and kept the dropout layers active to increase the model variability. We also employed three AF2 versions by using four seeds, while playing with the template usage and utilizing 3, 9, 21 recycles, while adjusting MSA pairing (Table 1). As an outcome, we run 11 different scenarios with ColabFold [10] (Methods).

**Table 1.** Our MinnieFold scheme.

| Scenario | AF2 version | Dropout | Templates | No of Recycles | MSA Pairing | No seeds x weights |
| --- | --- | --- | --- | --- | --- | --- |
| 1 | v2.1 | Yes | Yes | 3 | No | 4x5 |
| 2 | v2.2 | Yes | Yes | 3 | No | 4x5 |
| 3 | v2.3 | Yes | Yes | 3 | No | 4x5 |
| 4 | v2.1 | Yes | No | 3 | No | 4x5 |
| 5 | v2.2 | Yes | No | 3 | No | 4x5 |
| 6 | v2.3 | Yes | No | 3 | No | 4x5 |
| 7 | v2.1 | Yes | No | 21 | No | 4x5 |
| 8 | v2.2 | Yes | No | 9 | No | 4x5 |
| 9 | v2.3 | Yes | No | 9 | No | 4x5 |
| 10 | v2.3 | Yes | No | 21 | No | 4x5 |
| 11 | v2.1 | Yes | No | 21 | Yes | 4x5 |
| Total number of models generated |  |  |  |  |  | 220 |

In ranking these models, we focused on the average peptide pLDDT (pep-pLDDT) instead of the regular AF2 confidence score. This was done since we observed overinflated ipTM and pTM values due to the domination of off-interface antibody regions. The top pep-pLDDT (third ranking AF2 model) was generated by the 11^th^ scenario, which used AFv2.1 with 21 recycles, with no template information and paired MSA (Table 1). In the order of decreasing AF2 confidence scores, the pep-pLDDT scores of the top three models were obtained as 55, 62, and 68. Of note, all these models had almost identical ipTM values (Fig. 1A-C).

**Figure 1.**
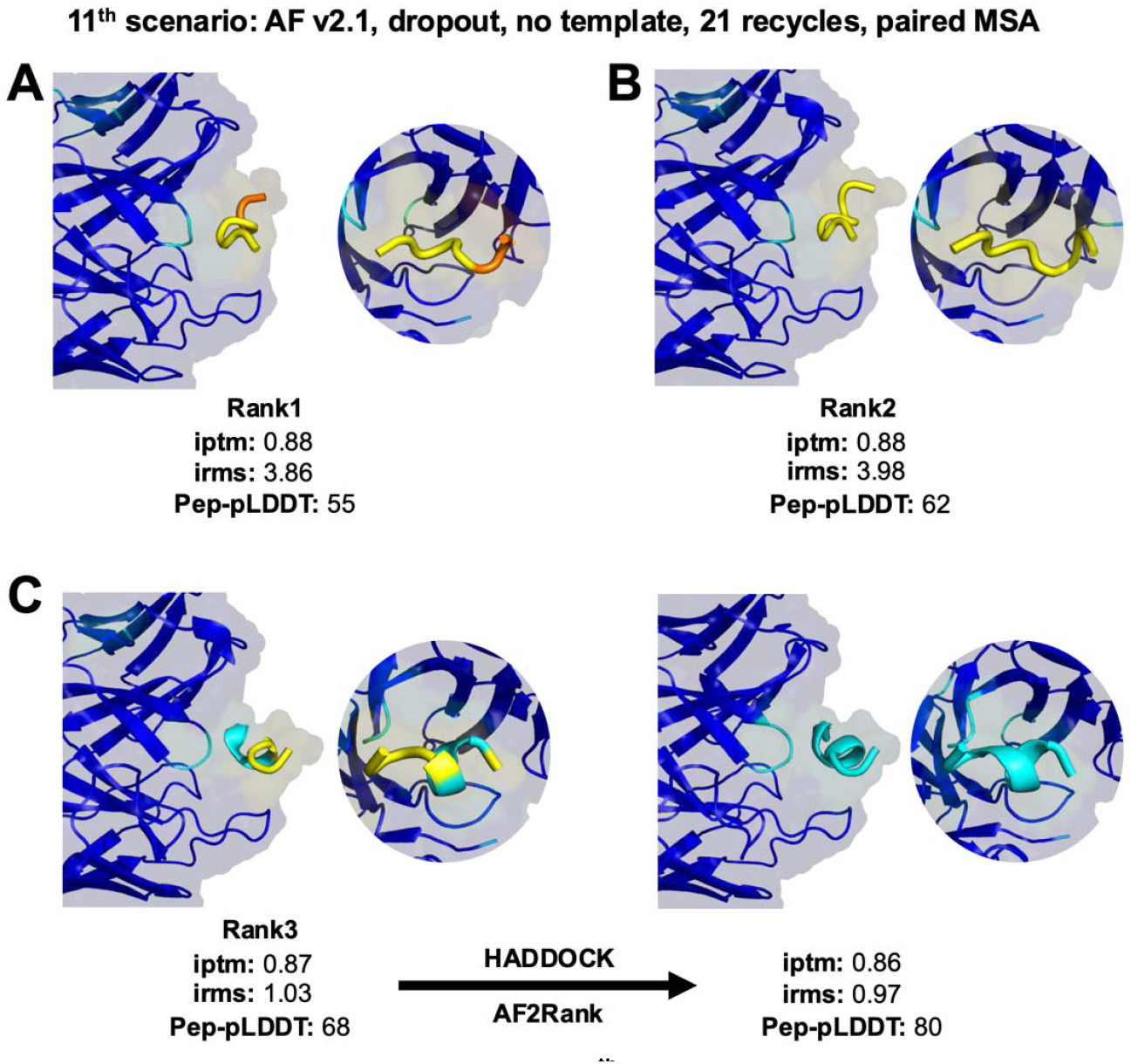
Evaluation of the top models from the 11^th^ MinnieFold scenario. **A**. The first, **B**. second, and **C**. third ranking models of the 11^th^ scenario. The change in third ranking model is represented by the arrow after refinement and rescoring (Table S1). All the models are colored according to their pLDDT scores and depicted in cartoon and transparent surface representation (pLDDT coloring: blue (highest), cyan, yellow, orange (lowest) confidence). ipTM, irms, and pep-pLDDT values of each model are provided below their associated structures.

Our top pep-pLDDT model was then subjected to HADDOCK 2.4 refinement [12] (since the relaxation module of ColabFold was down at the time of competition [22]). The final pep-pLDDT value of our HADDOCK-refined model was acquired through AF2Rank (Methods). AF2rank is a protocol developed to calculate the AF2 scoring metrics of a given structure [13]. It uses the input structure as a template and passes its sequence through AF2, while muting the input MSA information to prohibit further sampling. As an outcome, we acquired a pep-pLDDT of 80 and submitted this model as our top-ranking prediction (Fig. 1D). This model was classified as medium, placing us among the top seven predictors (Kozakov/Vajda, Pierce, Karaca, Huang, Venclovas, Zou, and Vakser). Our post-CAPRI analysis showed that the first two ranking models of the 11^th^ scenario were incorrect, while the third ranking one was acceptable. So, our submission became a medium model after AF2Rank scoring (Table S1). For comparison, we also relaxed the third ranking structure with AF2’s refinement protocol, which led to the same outcome as the HADDOCK refinement (Table S1).

Our other nine submissions came from a 450 ns long MD trajectory, which we started from a structure obtained through a default ColabFold-AFv2.3 run (Methods). At this point, we used MD, since we hypothesized that MD could aid us in finding a stable peptide pose. Accordingly, we saved a snapshot at each nanosecond and analyzed 450 conformers. Our submissions were selected according to their peptide-antibody contact profiles (the higher interface contacts were ranked higher, Figure S1). All these models were classified as acceptable, except for the last incorrect one.

### MinnieFold sampling could explore a conformational space range up to 12 Å for T232

T232 was made of regulatory and catalytic subunits of a serine/threonine-protein phosphatase 2A (PP2A) in complex with a TIP41-like (TIPRL) protein. In an available complex structure, TIPRL was found to interact with both PP2A subunits (Fig. 2A, PDB ID: 5W0W [23]). The main challenge in this target was to identify an alternative heterotrimeric organization of this complex. To find this conformation, we employed our MinnieFold sampling while skipping 9^th^, and 10^th^ scenarios to save time (since ColabFold was down at the time of competition [22]) (Table 1). Among the generated conformations, we wanted to submit models with a wide RMSD range to increase our chances in capturing the alternative binding mode. Our modeling scenarios resulted in Cα−RMSD values ranging up to 11.7 Å (Fig. 2B), primarily due to the clockwise movement of TIPRL (Fig. 2C). The models with the highest conformational variability came from the 8^th^ scenario (Table 1) (Fig. 2C). Based on the observed RMSD variation, we manually selected our first eight submissions, all of which are classified as medium (Figure S1).

**Figure 2.**
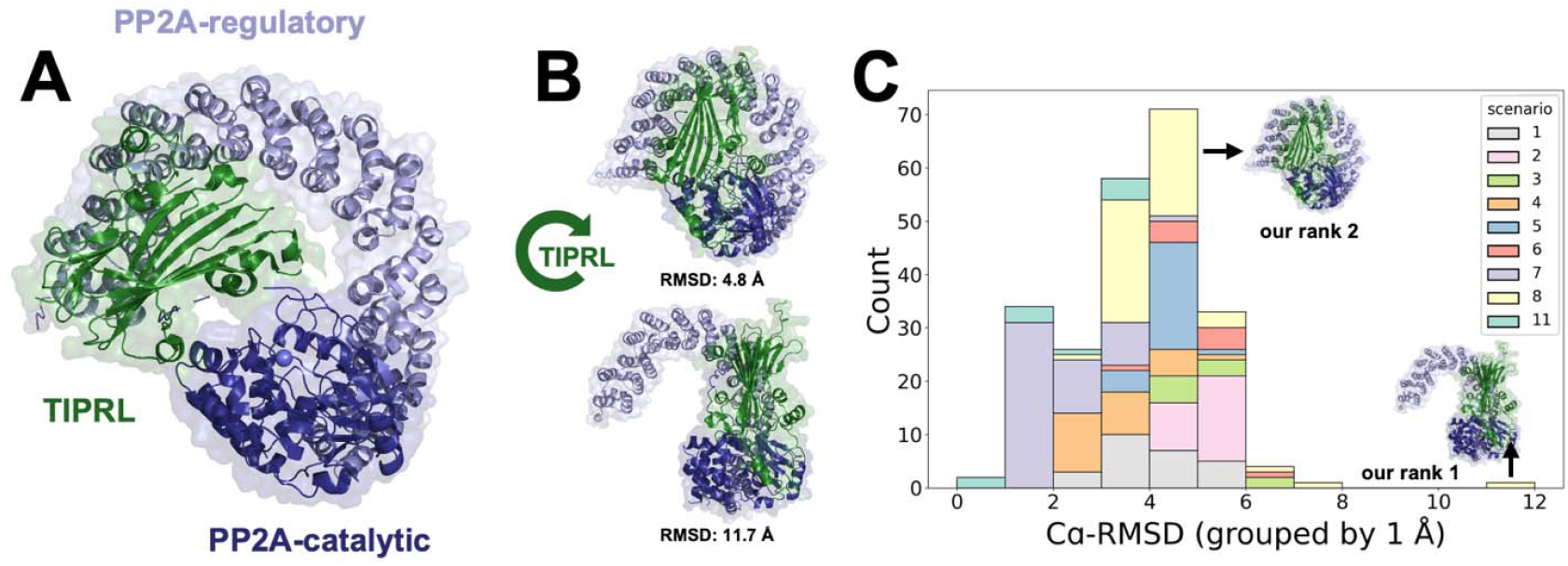
Structural comparison of T232 models to the reference structure (PDB ID: 5W0W [20]). **A**. TIPRL in complex with the regulatory and catalytic subunits of PP2A. **B**. Structural depiction and the RMSD values of our top two submissions RMSD of 11.7 Å and 4.8 Å. **C**. The bar graph of Cα-RMSD values across all generated models colored according to their scenarios. The reference state was chosen as the top-ranking structure of the 11^st^ scenario. Our top two submissions are highlighted on the plot.

As a secondary approach, we aimed to generate a conformationally diverse set of models using HADDOCK in the *ab initio* mode. All models generated in it1 and water stages converged to the arrangement presented in 5W0W. We, therefore, selected our last two submissions from it0 manually from the high Cα-RMSD regime. Unfortunately, these models turned out to be incorrect.

### MinnieFold could have prevented our failure in T233, but not in T234

T233 and T234 were about predicting two different antibody binding sites on a major histocompatibility class (MHC) I complex. MHC type I complex structures are essential for the immune system, where their HLA-A component recognizes an 8-10 amino acids long antigenic peptide (Fig. 3A) [24]. In the absence of this peptide, the peptide-binding site of MHC is stabilized with a tapasin (TAP) molecule [25]. Since, for both targets, the presence of an antigenic peptide was not reported, we assumed that an antibody could bind on HLA-A either on its peptide binding site (Ab(1) in Fig. 3A), its TAP binding site (Ab(2) in Fig. 3A) or the opposite of its TAP binding site (Ab(3) in Fig. 3A). Based on this assumption, we devised three data-driven docking scenarios, where we inferred the HLA-A binding sites from previously published complexes (PDB ID: 1QVO [26] for Ab(1) and PDB ID: 5WER [27] for Ab(2) and Ab(3)). Data-driven docking was carried out with HADDOCK. As the interface sizes of each site differed, we used FoldX to rank the HADDOCK-generated structures. We made sure that our submitted models contained at least one solution from each targeted binding site. Unfortunately, this strategy did not lead to an acceptable solution.

**Figure 3.**
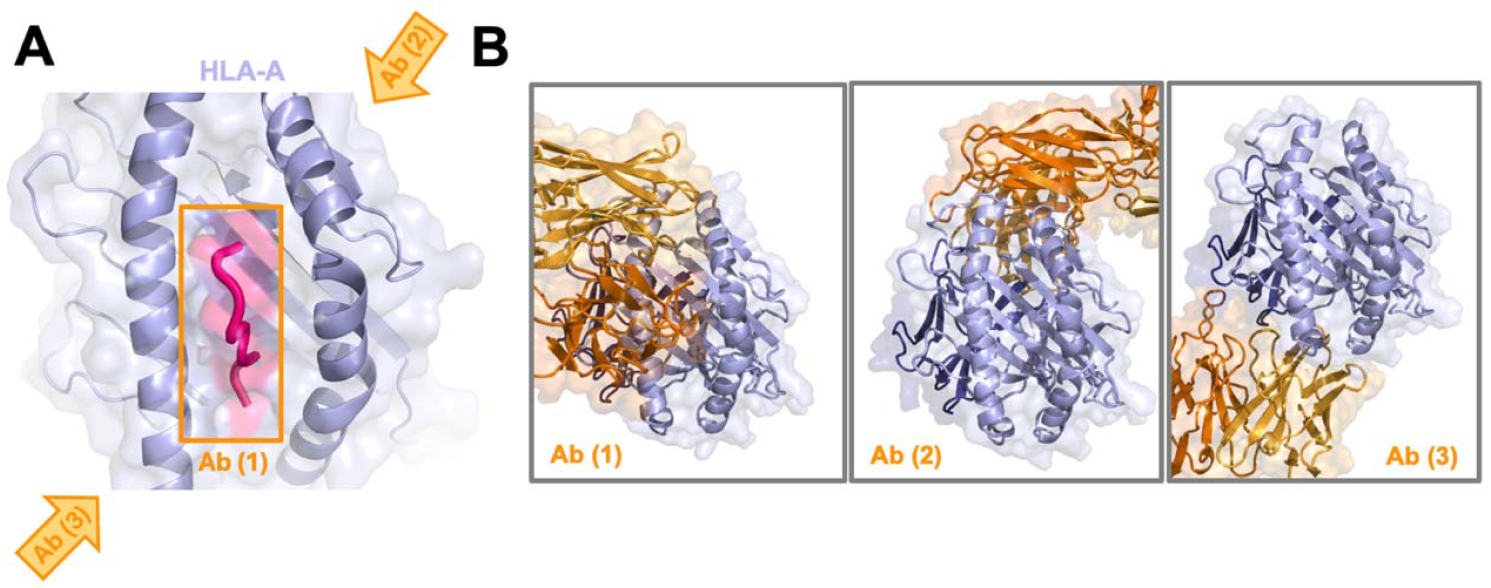
The antibody (Ab) binding sites used in the modeling of T233 and T234. **A**. The HLA-A and antigenic peptide binding shown in top view. The targeted Ab binding sites are shown by orange arrows and a box on the reference structure. **B**. Examples from the generated models for each targeted Ab binding site. MHC-I and antibodies are represented in lilac and orange cartoon and shades, respectively.

As post-CAPRI analysis, we ran our MinnieFold sampling on T233 and T234. Due to the large size of these targets, we reduced the number of seeds to two for each modeling, leading to 110 models per target. For T233, even with this reduced MinnieFold protocol, we could obtain one medium and one acceptable solution. The medium-quality model came from the 2^nd^ scenario, while the acceptable model came from the 1^st^ (Table 1, Table S1). Interestingly, the medium model had the highest confidence score among all models, while the acceptable one was ranked at 85. So, by using AF v2.2 with 3 recycles and template information enabled, we could have submitted a top-ranking medium prediction. For T234, on the other hand, our protocol could not generate a meaningful solution. For a final analysis, we applied our refinement and rescoring protocol to the acceptable and medium quality models of T233. As a result, our acceptable model improved to medium quality, with a notable increase in the DockQ score from 0.4 to 0.7 (Table S1). So, as claimed in the AF2Rank paper, it is indeed possible that AF2Rank can push an energetically favorable (acceptable) model downhill to an even more energetically favorable state (medium) [15].

### Scoring challenge

For the scoring challenge, we scored the chain-mapped interfaces with the *AnalyseComplex* function of FoldX (Methods) [9]. We then visually inspected the top 100 models with the lowest energies using PyMOL 2.5.2 to make the final selection [19]. This strategy helped us to select two acceptable models for T231, though they were our 4^th^ and 9^th^ ranking submissions. For T232, we scored eight medium-classified models out of our ten scoring submissions, while having our 1^st^ and 5^th^ submissions as incorrect. In Figure 4, we shared the FoldX energy distribution over the entire scoring set and highlighted the selected energy range for our submissions. Except for T232, our selections did not explore the lowest energy values, which may have contributed to our lower performance in these cases. Considering the FoldX energy distribution and our performance, it is clear that our manual inspection had a major diminishing effect on our performance.

**Figure 4.**
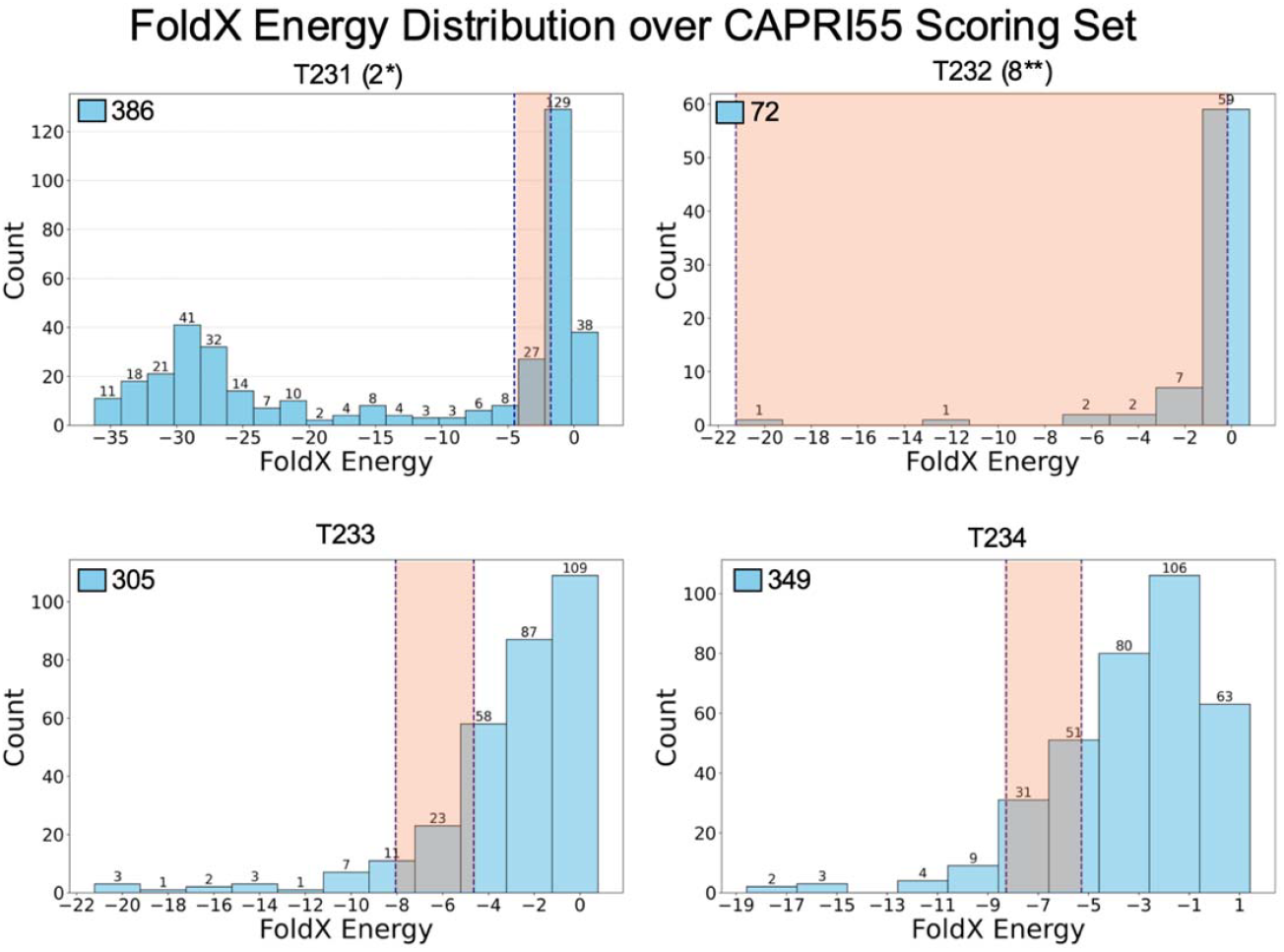
FoldX Energy Distribution for the CAPRI55 Scoring Set. Only models with negative energies are represented. The light orange shaded panels indicate the energy range of spanned by our submissions for the scoring challenge. We specified our performance with the number of picked medium (**), and acceptable (*) quality models, if available.

## DISCUSSION

In this work, we demonstrated that a 50-fold reduced version of Wallner’s massive sampling, MinnieFold, could generate medium quality models for T231, T232, and T233. For ranking T231, though, replacing the standard AF2 scoring was needed since ipTM and pTM values of this target was dominated by the high confidence off-interface antibody regions. Notably, others have also identified limitations in AF2’s native scoring for antibody-antigen complexes and proposed alternative metrics, such as interface-based scoring strategies, to improve performance [28]. Following these insights, we applied a peptide-specific interface-based metric, pep-pLDDT, to accurately score antibody-peptide complexes. This targeted adjustment underscores the necessity of tailored scoring metrics for specialized antigen-antibody systems to achieve reliable model evaluations.

We further showed that employing MinnieFold sampling could generate a wide range of conformations in the case of T232, demonstrating another application area of our protocol. Importantly, incomplete information on T233 and T234 made us design targeted docking strategies, which did not result in a successful model generation. Interestingly, a simple AF2v2.2 run could have produced a medium solution for T233. It is worth noting that, in this study, we exclusively used ColabFold, which relies on MMseqs2 for constructing MSAs, as opposed to DeepMind’s AlphaFold pipeline, which uses JackHMMer and HHblits. This choice reflects the accessibility and computational efficiency of ColabFold but also may introduce differences in the quality of MSAs, which can impact the model quality. We also used dropout enabled in all scenarios to enhance model diversity.

In CAPRI55, another team, the Brysbaert Team, also followed a Wallner-like sampling through their MassiveFold tool [5, 29]. This approach uses a GPU parallelization pipeline to significantly reduce the time required to run AFSample by using all available AF2 versions. As an outcome, MassiveFold generated 6,030 models per target. This strategy led to medium for T231, high for T233, and acceptable models for T234 [30]. Interestingly, AF2 scoring could not pick most of these solutions. So, generating more models in such co-evolution independent cases apparently makes even more problematic.

After CAPRI55, DeepMind released AlphaFold3 (AF3), making it accessible to researchers via a web server [11]. AF2 was based on a transformer based Evoformer architecture, leveraging multiple sequence alignments (MSAs) and attention-based pair representations for structure prediction. AF3 replaces Evoformer with Pairformer module simplifying MSA processing. Furthermore, AF3 introduces a diffusion-based generative model. Given its unique architecture, we assessed AF3’s performance on T231 and T233-4 antibody targets by using a single and five seeds (recommended for multimer predictions). Notably, AF3 failed to generate an acceptable model for T231 (Table S1). This may be attributed to the fact that the original AF3 study utilized 1000 seeds in antibody-antigen modeling [13]. For T232, using five seeds did not result in diverse conformational generation (mean backbone RMSD 1.25 ± 0.43 Å). In contrast, AF3 successfully produced high-quality models for T233, surpassing the performance of AF2 (Table S1). However, T234 still presented a challenge, as AF3 attempted to fill the peptide-binding site with random protein segments. This suggests that T234 is particularly difficult to model in the absence of the peptide. Based on these results, for antibody-antigen targets, we recommend the combined use of enhanced AF2 and AF3 sampling.

To conclude, CAPRI55 provided an excellent platform to test our MinnieFold sampling approach. The accuracy achieved was better than we had anticipated, demonstrating that this method could serve as a green, user-friendly option for antibody complex modeling where the typical co-evolution signal is missing. While the small number of targets in CAPRI55 limits a comprehensive evaluation, the promising results highlight the potential of MinnieFold as a computationally efficient and accessible approach, particularly through ColabFold’s web service. Our MinnieFold strategy significantly reduced GPU hours and, consequently, energy consumption by 95%. For perspective, Wallner’s Team used 41,200 GPU hours to model 38 targets in CASP15-CAPRI, this computational effort requires around 397 trees a full year to offset the carbon footprint (12,360 kWh, 8.74 tons of CO_2_ emissions) [31]. Thus, with its reduced computational cost, MinnieFold holds great promise for widespread use and future applications.

## DECLARATIONS

### Data Availability Statement

The data that support the findings of this study are available from GitHub at https://github.com/CSB-KaracaLab/MinnieFold.

### Author Contributions Statement

**Büşra Savaş**: Conceptualization; Investigation; Writing - original draft; Methodology; Visualization; Writing - review & editing; Formal analysis.

**İrem Yılmazbilek**: Investigation; Formal analysis; Methodology; Writing - original draft.

**Atakan Özsan**: Investigation; Formal analysis; Methodology.

**Ezgi Karaca**: Writing - original draft; Writing - review & editing; Project administration; Supervision; Conceptualization; Investigation; Methodology; Funding acquisition.

All authors reviewed and approved the final manuscript.

