## Supporting Information for "Towards a greener AlphaFold2 protocol for antibody-antigen modeling: Insights from CAPRI Round 55"

**Running Title:** MinnieFold: Green Enhanced AF2 Sampling

Büşra Savaş^1,2^, İrem Yılmazbilek^1^, Atakan Özsan^1^, Ezgi Karaca^1,2^

^1^İzmir Biomedicine and Genome Center, İzmir, Türkiye;

^2^Izmir International Biomedicine and Genome Institute, Dokuz Eylül University, Izmir, Türkiye

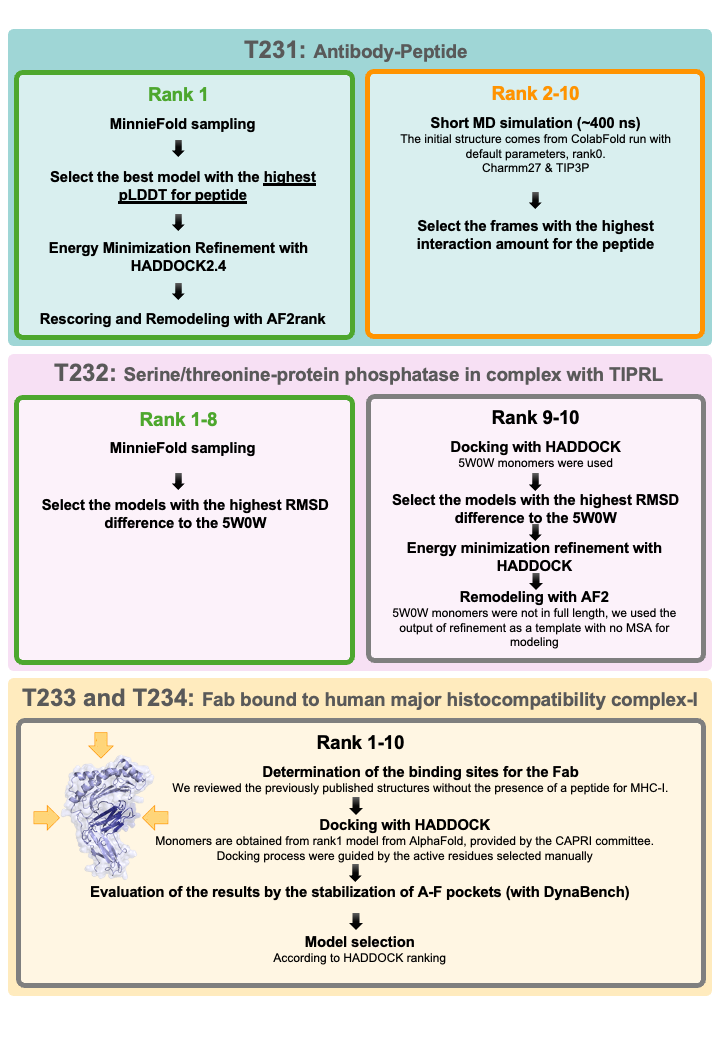

**Figure S1. Our case specific CAPRI55 modeling approaches.** Frame colors indicate our performance with the modeling approach: green (medium), orange (acceptable), and gray (incorrect).

**Table S1.** CAPRI classification for the noted antibody-antigen models of T231, T233, and T234. All CAPRI metrics are calculated with respect to the native structures.

| **ID** | **dockq** | **irms** | **lrms** | **fnat** | **classification** |
| --- | --- | --- | --- | --- | --- |
| T231-11^th^scenario_rank1 (Figure 1A) | 0.276 | 3.864 | 7.444 | 0.130 | Incorrect |
| T231-11^th^scenario_rank2 (Figure 1B) | 0.284 | 3.979 | 7.639 | 0.174 | Incorrect |
| T231-11^th^scenario_rank3 (Figure 1C) | 0.623 | 1.023 | 6.027 | 0.522 | Acceptable |
| T231-11^th^scenario_rank3 + Rosetta Refinement | 0.621 | 1.045 | 6.007 | 0.522 | Acceptable |
| T231-11^th^scenario_rank3 after HADDOCK | 0.653 | 1.016 | 6.040 | 0.609 | Acceptable |
| T231- 11^th^scenario_rank3 after HADDOCK + AF2Rank  our-top-model  (Figure 1D) | 0.702 | 0.9687 | 5.494 | 0.696 | Medium |
| T231-AF3_rank1 | 0.345 | 3.646 | 7.151 | 0.304 | Incorrect |
| T233-2^nd^scenario_rank1 | 0.760 | 1.076 | 3.949 | 0.796 | Medium |
| T233-2^nd^scenario_rank1 + Rosetta Refinement | 0.751 | 1.091 | 3.940 | 0.776 | Medium |
| T233-2^nd^scenario_rank1 after HADDOCK | 0.744 | 1.133 | 3.969 | 0.776 | Medium |
| T233-2^nd^scenario_rank1 after HADDOCK + AF2Rank | 0.760 | 1.141 | 4.129 | 0.837 | Medium |
| T233-1^st^scenario_rank1 | 0.453 | 3.009 | 7.111 | 0.571 | Acceptable |
| T233-1^st^scenario_rank1 + Rosetta Refinement | 0.405 | 3.007 | 7.116 | 0.429 | Acceptable |
| T233-1^st^scenario_rank1 after HADDOCK | 0.393 | 3.137 | 7.149 | 0.408 | Acceptable |
| T233-1^st^scenario_rank1 after HADDOCK + AF2rank | 0.711 | 1.497 | 4.229 | 0.837 | Medium |
| T233-AF3_rank1 | 0.726 | 0.941 | 6.621 | 0.837 | High |
| T234-AF3_rank1 | 0.034 | 10.702 | 28.316 | 0 | Incorrect |
